# Meltome Atlas of *Arabidopsis thaliana* Proteome: A Melting Temperature-Based Identification of Heat & Cold Resistant Proteins

**DOI:** 10.1101/2025.02.22.639691

**Authors:** Karan Martens Mohanta, Tapan Kumar Mohanta

## Abstract

The study elucidated the thermal properties of the entire *Arabidopsis thaliana* proteome by considering the melting temperature (Tm) and the melting temperature index (TI). In total, 48359 protein sequences were analyzed, and the melting temperature of the proteins was recorded in three groups (Tm < 55°C, 55-65°C, and > 65°C). The Tm index of the *A. thaliana* proteome ranged from -15.6008 (< 55°C) to 9.605 (> 65°C). At least 22826 proteins were found in the Tm group of 55°C to 65°C, 20640 proteins were found in the Tm group of > 65°C, and only 4893 proteins were found in the Tm group of < 55°C. The mediator of RNA polymerase II transcription subunit-like protein was found to possess the highest Tm index (9.60), while the NADH dehydrogenase 5B subunit was found to contain the lowest TI (-15.60). The amino acid composition analysis of the *A. thaliana* proteome revealed that the frequency of Ala, Asp, Glu, Gly, Lys, Gln, and Val increased with the increase in Tm, while the amino acids Cys, Phe, and Trp decreased with the increase in the Tm of the *A. thaliana* proteome. The molecular mass of the *A. thaliana* proteome ranged from 0.149 to 611.888 kDa, and protein in the Tm group at 55-65°C showed the highest average molecular mass. The machine learning analysis revealed an increase in the molecular mass positively correlated with the increase in the Tm of the proteins. The codon usage pattern revealed, the codon pair prefer the Tm group specific occurrence where ATG-ATG, CAA-CAA codon pairs were predominated. Relative synonymous codon usage of the three Tm groups revealed AGA (Arg) and CCA (Pro) were the preferred codons for the low and high Tm group DNA sequences, respectively. Codon context analysis revealed the presence of preferences of the Tm group specific codon pairing. There was a variation in the nucleotide position of the codons in different Tm groups. Evolutionary study revealed, gene duplication was the predominant evolutionary feature and all of the studied genes in the three Tm group undergone duplication. Codon context analysis revealed distinct clustering pattern in high Tm protein group. The study underscores the role of amino acid composition, molecular mass, and codon usage in determining the thermal stability of the proteins in the *A. thaliana*. The study reflected the evolution of high Tm-adapting genes through gene duplication, highlighting the role of gene and genome evolution towards encoding high Tm proteins for stress resilience.

## Background

Plants are ubiquitous sessile organisms, and hence they face a plethora of stresses and environmental challenges. These challenges adversely affect the growth, development, and productivity of the plants [1, 2]. However, they have established a tremendous ability to respond and adapt to these stresses for their survival [3, 4]. The translation product, proteins, is crucial to these events, and they play a fundamental role in these stress responses [5–7]. They act as key signaling molecules and key regulators of plant defence responses [8–10]. They perceive, respond, regulate, and adapt to various stress conditions. The roles of receptor-like kinase (RLK) and receptor-like proteins (RLP) are tremendous in perceiving external stress signals [8, 11–13]. Some proteins act as second messengers like calcium-binding proteins (calcium-dependent protein kinase, calmodulin, calmodulin-like protein, calcineurin B-like proteins) [14–18], and phospholipases [19,20]. Transcription factors like NAC, WRKY, MYB, AP2/ERF, and bZIP regulate the expression of several stress-responsive genes in plants [21–24]. These TFs bind to the specific promoter region and regulate the gene expression, thus adapting to the stress responses [25, 26]. Kinases (MAPKs) and phosphatases (PP2C) play phosphorylation and dephosphorylation events to alter the protein function [27, 28]. Ubiquitin ligase conducts ubiquitination through the ubiquitin-proteasome complex and removes the damaged proteins and thus regulates the function of key stress proteins [29,30]. During the stress event, the generation of reactive oxygen species occurs, and the oxidative stress gets mitigated by superoxide dismutase (SOD), peroxidase (POD), and catalase (CAT) [31]. Similarly, osmotic adjustment in cells is regulated by proline dehydrogenase, trehalose 6-phosphate, and other proteins and balances the osmotic stress condition [32]. During heat stress, heat shock protein (HSP) helps in protein folding and prevents aggregation of denatured proteins resulting from the heat stress [33]. The role of HSP protein is extraordinary in maintaining protein homeostasis [34, 35]. Like HSP, aquaporin protein regulates drought stress in plants [36–38].

From several environmental stresses, heat and cold stress are the most common types of stresses. Cold stress includes chilling stress (0-15°C) and freezing stress (below 0°C) [39]. The chilling and freezing stress arrest the proteomic and enzymatic activities and thus slow down the metabolic process [39, 40]. The cold stress leads to the production of Osmo protectants like proline through the enzyme pyrroline-5-carboxylase synthetase (P5CS) [41–43]. Similarly, sucrose-phosphate synthetase (SPS) and fructan fructosyltransferase (FFT) help in sugar metabolism [44]. Unlike cold stress, when temperature rises above 35°C, plants encounter heat stress. The heat stress greatly impacts the protein stability and membrane permeability [45, 46]. The heat stress is sensed by protein denaturation and changes in membrane permeability [47, 48]. The HSPs help in protein folding and prevent the aggregation of denatured and damaged proteins [33, 49]. Sometimes plants encounter a combined stress response of heat and drought stress [50]. During this event, proteins associated with crosstalk get involved, and their signaling mechanism helps to mitigate the stresses. Thus, proteins are the pivotal elements involved in heat and cold stress mitigation by orchestrating complex networks of signalling pathways and metabolic adjustment [51, 52]. A lot of studies are being conducted to identify and implement different genes and proteins involved in such stress tolerance for the development of stress-tolerant crops. Indeed, this is also pivotal for the development of sustainable agriculture in the face of global warming. So that plants can thrive in challenging environmental conditions.

More specifically, when a plant undergoes extreme heat or chilling stress, the protein either undergoes denaturation or strong folding/misfolding [53,54], thus impacting the protein function. Proteins with a higher melting temperature tend to be more stable and can uphold their structure and function at the higher temperature [55–57]. This can be quite crucial to withstand the heat stress. Similarly, proteins with cold stress can also undergo denaturation (less common) and can lead to disruption in cellular and metabolic activities [58,59]. Cold stress can alter the secondary and tertiary structures of the protein, making them quite rigid and nonfunctional [60–62]. The proteins associated with the membrane and structural components of cells will be greatly impacted due to such chilling stress. Prolonged cold stress can lead to protein aggregation, leading to the formation of insoluble protein complexes and ultimately cellular death. Cold temperatures will slow down the enzyme activities and their kinetics, thus reducing their biochemical reactions. If any protein associated with the Calvin cycle gets impacted by chilling stress, there will be a hindrance in photosynthesis, leading to poor growth and development of the plants. Similarly, chilling stress can precipitate the membrane-associated lipid thus disrupting membrane fluidity and signaling pathways. Protein-protein interactions were also greatly impacted due to the denaturation of proteins, leading to alternations in charges and protein conformation. Further, post-translational modification gets impacted by such chilling or heat stress, which regulates protein activity and stability.

A lot of studies are conducted to find different genes and proteins associated with stress tolerance in plants [63, 64]. Starting from 2D protein gel electrophoresis to single-cell transcriptome and from small nucleotide polymorphism to marker-assisted identification of novel traits, several such approaches are being implemented to identify novel genetic traits in plants [65–67]. These approaches are still in force to identify novel traits in plants. However, so far, no study is being conducted to identify the novel genetic and genomic traits using the physiological parameters of their translated product, i.e., protein. Proteins are quite sensitive to pH change, which affect their structure, function, and physiochemical properties. Further, physiological parameters like the melting temperature (Tm) of proteins can be one of the most important factors that can play a crucial role in protein function. Tm is a temperature at which protein undergoes denaturation and loses its three-dimensional structure and function. At Tm, half of the protein remains in a folded (native) state, and half of the protein remains in an unfolded state. It is one of the important parameters to understand the protein stability. A higher Tm indicates higher stability of the protein, where the protein can maintain and sustain its three-dimensional shape and function. The amino acid composition of the protein greatly influences the thermal stability and overall structure of the protein [68, 69]. Further, larger and more complex proteins might have more intricate folding and may have a higher melting temperature. The presence of hydrophobic amino acids in the protein tends to increase the Tm of the protein and thus protein stability [70–72]. Similarly, a properly folded protein has a higher Tm compared to partially or misfolded proteins. Further, post-translational modifications like glycosylation and phosphorylation affect protein stability [73, 74] through its Tm. The presence of other factors like pH, ionic strength of the protein, and stabilizing agents can impact the Tm of the protein. The presence of acidic (aspartic acid and glutamic acid) and basic (lysine and arginine) ionization groups affects the electrostatic interactions of the proteins and stabilizes the protein structure [75,76]. This shows that Tm is a fundamental property of the protein that influences the protein’s stability, function, and interactions. Therefore, it becomes crucial to understand this parameter to understand the protein’s behavior under various environmental conditions. By analyzing Tm of proteins, we can understand the basic insight of the mechanism of the protein by which the protein adapts to the harsh environmental condition and mitigates stress. This physiochemical parameter of the protein can be quite crucial to identify the stress-tolerant proteins and subsequently stress-tolerant crops for biotechnological application. Therefore, we identified the Tm values of all the protein sequences of *Arabidopsis thaliana* proteome and reported them

## Materials and Methods

### Retrieval of Arabidopsis proteome sequences

The protein sequences of *Arabidopsis thaliana* were downloaded from the TAIR (The *Arabidopsis* Information Resources) database. The proteome file contained the translated protein sequences of all the coding DNA sequences (CDS) and their splice variants. It included a total of 48359 protein sequences. The downloaded protein sequences were subjected to analysis of melting temperature (Tm).

### Prediction of Melting Temperature

The melting temperature prediction of the entire *Arabidopsis thaliana* proteome was made using the “Melting temperature prediction” (http://tm.life.nthu.edu.tw/) pipeline [77]. The online platform calculates the melting temperature and melting temperature index (TI) of the protein. Only one sequence can be submitted at once to find the melting temperature and melting index. The resulted Tm and TI were recorded in a Microsoft Excel file for further analysis.

### Prediction of molecular mass and isoelectric point of Arabidopsis thaliana proteome

The molecular mass and isoelectric point of all the chloroplast protein sequences were calculated using the IPC-isoelectric point calculator (http://isoelectric.org/), version 2.0 [78]. The IPC Python software was downloaded and run on a Linux-based command-line platform. Command lines were used as per the instructions of the software. The IPC software resulted in the molecular mass and isoelectric point of individual proteins. We saved the data in text file format for further analysis.

### Statistical analysis of Arabidopsis thaliana Proteome Melting temperature

The melting temperatures of the *Arabidopsis thaliana* proteome were divided into three groups. They are as follows: (I) < 55°C, (II) 55-65°C, and (III) > 65°C. Later, the protein sequences of the *A. thaliana* proteome were segregated according to their melting temperature, and subsequent analysis was performed. Correlation analysis of proteins with Tm < 55°C, 55-60°C, and > 65°C with respect to molecular mass and isoelectric point was conducted using JASP 0.19.0.0 version software. Pearson’s correlation with a 95% confidence interval was used for the study. One sample student t-test of the Tm index of *the A. thaliana* proteome was conducted using JASP 0.19.0.0 version software to confirm that the mean is different from zero (*p* < 0.01). A frequency distribution of molecular mass and isoelectric point of all three Tm groups was conducted with *p* < 0.05 using Past4 software. The amino acid composition of *the A. thaliana* proteome was calculated based on their melting temperature groups by running a Linux-based command. All other basic statistical details were calculated using Microsoft Excel 2016.

### Tm Based Evolutionary Analysis of A. thaliana proteome

To understand the evolutionary details of the *A. thaliana* proteomes, we selected the top 50 protein sequences from three TI groups. Group I contained the bottom 50 TI protein sequences (< 55°C), group II contained the top 50 TI protein sequences of the 55-65°C temperature range, and group III contained the top 50 TI protein sequences with Tm > 65°C. The CDS sequences of the *A. thaliana* proteins were retrieved from the TAIR database, and a multiple sequence alignment was conducted within the individual TI group using MAFT software [79]. The multiple sequence alignment of the protein sequences was saved in CLUSTALW file format that was subsequently converted to .AL file format using MEGA7 software [80]. The resulting .aln file was subjected to model selection to conduct the phylogenetic analysis in MEGA7 software [80]. Later, we subjected the .aln file to phylogenetic analysis. Based on the resulting models, the best model was used to construct the phylogenetic tree. Various statistical parameters used to construct the phylogenetic tree were as follows: statistical method, maximum likelihood, test of phylogeny; bootstrap method, model/method, Tamura-Nei model. Gamma distribution with discrete gamma categories (5) was used to construct the phylogenetic tree. The nearest-neighbour-interchange (NII) ML heuristic method was used with a strong branch swap filter. There were no gaps or missing data treatment, and all sites were used to construct the phylogenetic tree. The resulting phylogenetic tree was saved in Newick file format and referred to as a gene tree.

To understand the gene loss, duplication, and divergence, it was important to compare the respective gene tree with their species tree. The species tree was constructed using the following link: https://www.ncbi.nlm.nih.gov/Taxonomy/CommonTree/wwwcmt.cgi. The gene tree and species tree were loaded and later reconciled in Notung software [81] version 2.9.1.5 to get the gene loss, duplication, and divergence. Relative synonymous codon usage and nucleotide position of codons were calculated using MEGA software version 7 [80].

## Results

### The Melting Temperature Index (TI) of Arabidopsis thaliana proteome ranged from 9.605 to -15.6008

The study was conducted to deduce the melting temperature and melting temperature index of the *Arabidopsis thaliana* proteome. All the protein sequences of the *A. thaliana* proteome were downloaded from the TAIR database. The proteome file included all the protein sequences, including the splice variants. Individual protein sequences were subjected to Tm analysis using the Tm predictor. The resulting Tm and TI were documented in an Excel file. It was found that the TI of the *A. thaliana* proteome ranged from -15.6008 (Tm < 55°C) to 9.605 (Tm > 65°C) (Table 1). The mediator of RNA polymerase II transcription subunit-like protein (At1g55080.2) was found to encode the highest TI (9.605), and the protein NADH dehydrogenase 5B subunit (AtMG00665.1) encoded the lowest TI (-15.6008) (Table 1). The melting temperature index was grouped into three categories (< 55°C, 55-65°C, & > 65°C) as mentioned above. It was found that the average TI of the *A. thaliana* proteome in the group of < 55°C was -0.623, for the group 55-65°C was 0.596, and for the group > 65°C it was 1.508. In the entire proteome of *A. thaliana*, there were 48359 protein sequences. From them, 20640 were found to contain Tm > 65°C, 22826 were found to contain Tm 55-65°C, and 4893 were found to contain Tm < 55°C. The average TI of the entire *A. thaliana* proteome was found to be 0.493.

### Trp was the lowest encoding amino acid in A. thaliana proteome with Tm > 65° C

In total, 48359 protein sequences of *A. thaliana* were found to encode 20855719 amino acids. From them, 1115151 amino acids were from the group with Tm < 55°C, 10696941 amino acids were from the group with Tm 55-65° C, and 9043627 amino acids were from the group with Tm > 65°C. It was found that the percentage of amino acid composition of Trp (1.135%) was lowest in the protein sequences that encode proteins with Tm > 65°C. The rate of Trp composition was highest (1.377%) in the proteins with Tm < 55°C. However, the percentage composition of Leu amino acid was highest (9.682%) in Tm 55-65°C and lowest (8.961%) in Tm < 55°C (Table 2). The percentage of amino acid composition of Ala, Asp, Glu, Gly, Lys, Gln, and Val was found to increase with the increase in the Tm of the proteins (Table 2, Figure 1). While the percentage of amino acid composition of Cys, Phe, His, Ile, Met, Asn, Arg, Ser, Thr, Trp, and Tyr was found to decrease with the increase in Tm of the proteins (Table 2, Figure 1). However, the composition of Leu amino acid was increased from Tm < 55 °C to 55-65 °C and decreased for Tm > 65 °C (Table 2).

**Figure 1.**
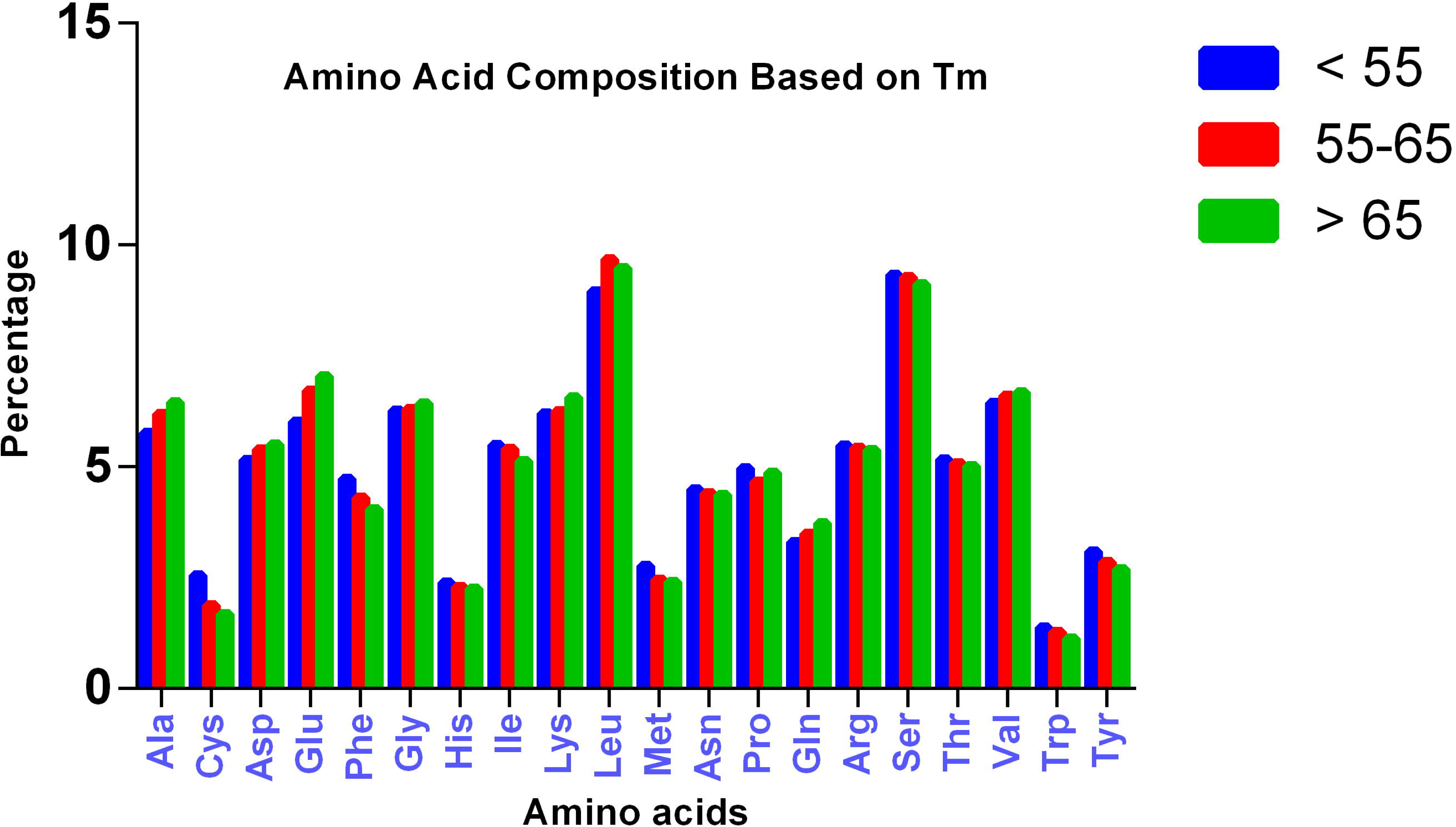
Amino acid composition of *Arabidopsis thaliana* proteome based on three different melting temperature range (< 55°C, 55-65°C, and > 65°C) of proteins. Cys and Trp were the lowest abundant amino acids found in Tm group > 65° C.

### The molecular mass of A. thaliana proteome ranged from 0.149 kDa to 611.888 kDa

It was pertinent to understand whether molecular mass plays any role in deciding the Tm of the proteins. Therefore, we calculated the molecular mass of individual proteins of the *A. thaliana*. It was found that the *A. thaliana* proteome encodes proteins from 0.149 kDa to 611.888 kDa. The average molecular mass of proteins of the Tm group > 55°C, 55-65°C, and > 65°C was 25.617, 52.482, and 48.924 kDa, respectively (Figure 2). The molecular mass of proteins with Tm group 55-65°C was the highest, whereas the proteins with Tm group < 55°C was the lowest (Figure 2). The protein that encoded the lowest molecular mass (0.149 kDa) protein in *A. thaliana* proteome was a hypothetical protein (AT1G64633.1) with a Tm of 55-65°C and a TI index of zero. Similarly, the highest molecular mass protein of the *A. thaliana* (611.888 kDa) proteome was a midasin-like protein (AT1G67120.2). It has a Tm of > 65°C with a TI index of 1.223. A correlation analysis of the molecular mass of *A. thaliana* proteome with three different Tm groups (< 55°C, 55-65°C, and > 65°C) was conducted to understand if the Tm groups with respect to molecular mass are correlated (Figure 3). The correlation plot was < 55°C vs. 55-65°C, < 55°C vs. > 65°C, and 55°C-65°C vs. > 65°C with an *r* value of 0.044, 0.002, and 0.021 (Figure 3). The frequency distribution of the molecular mass of *A. thaliana* proteome is depicted in Figure 2.

**Figure 2.**
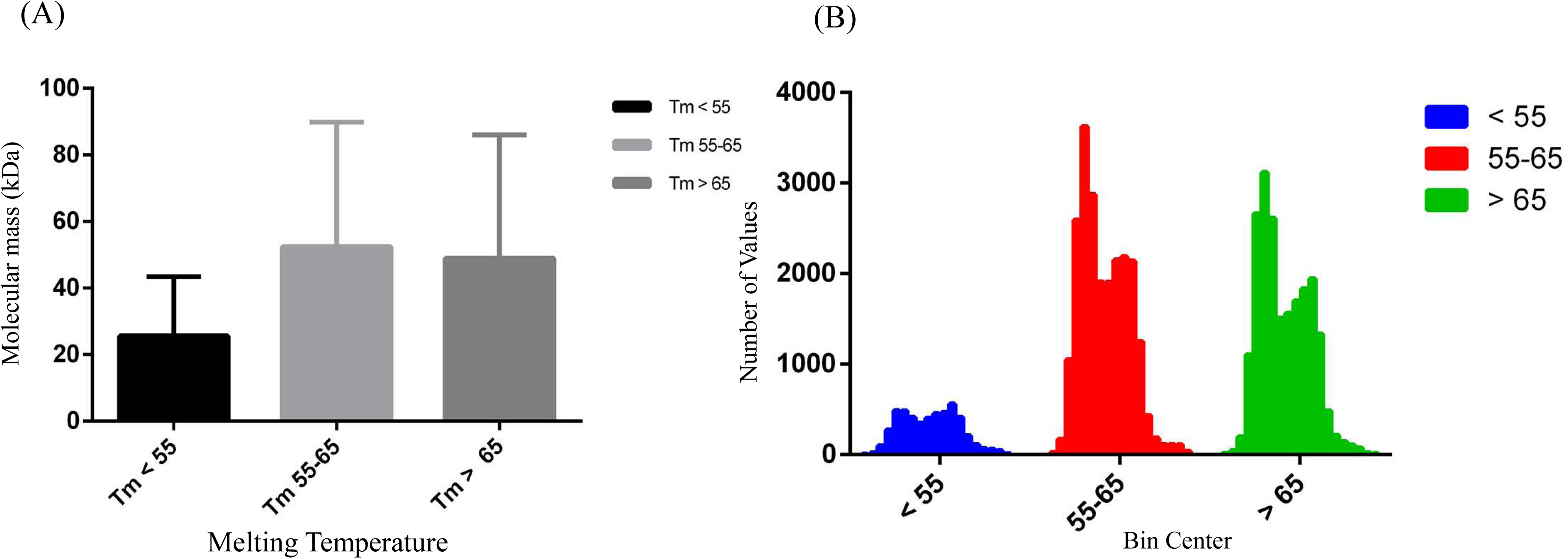
Figure depicting the (A) molecular mass (kDa) and (B) frequency of occurrence of the molecular mass of *A. thaliana* proteome in different Tm groups. The molecular mass of the proteins with the lowest Tm group < 55°C was found to be the lowest while the molecular mass of proteins with Tm > 65°C was moderate, and the molecular mass of proteins in Tm group 55-65°C was highest.

**Figure 3.**
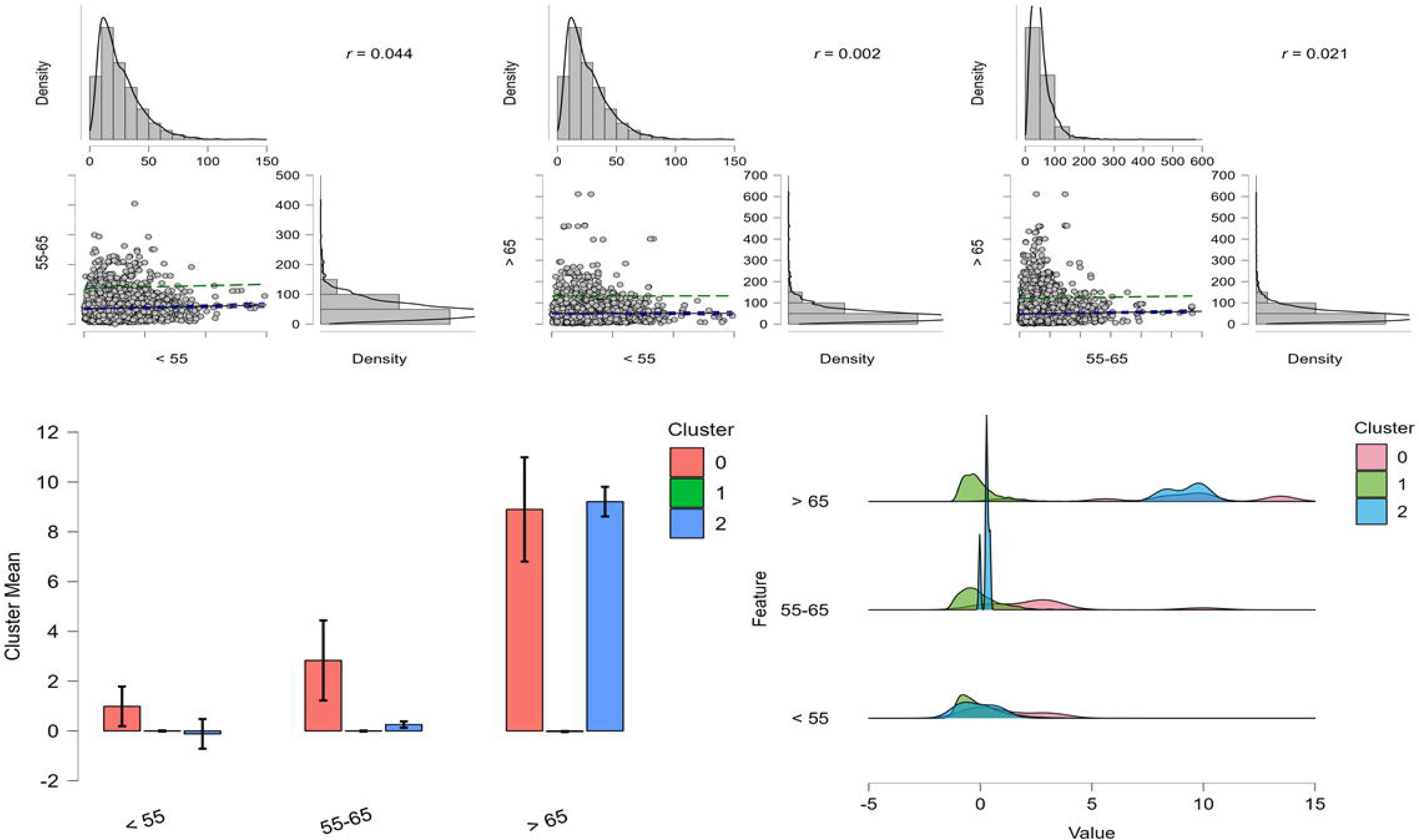
Correlation analysis between molecular mass (kDa) of different Tm groups. The correlation was studied between the molecular masses of Tm group of 55-65°C vs < 55°C, > 65°C vs < 55°C, and > 65°C vs 55-65°C. Correlation coefficient of molecular masses between the Tm groups 55-65°C vs < 55°C was found highest (*r* = 0.044). however, the correlation coefficient of molecular masses between the Tm groups was not significant to draw any conclusion. Pearson’s correlation was used to conduct the analysis with p < 0.05. The statistical analysis was conducted using the software JASP version 0.19.0.0.

### Isoelectric point (pI) of A. thaliana proteome ranged from 2.753 to 12.749

The isoelectric point of the *A. thaliana* proteome was calculated to understand whether the isoelectric point of the protein has any role towards the Tm of proteins. It was found that glycine-rich protein (AT3G44950.1) encoded the lowest *pI* (2.753) and ribosomal protein L41 family (AT3G56020.1) encoded the highest *pI* (12.749). The average *pI* of the *A. thaliana* proteome was 6.78. The average *pI* of the *A. thaliana* proteome with Tm < 55°C was 7.11; the *pI* of proteins with Tm 55-65°C was 6.75, and the *pI* of proteins with Tm > 65°C was 6.74 (Figure 4). A correlation analysis was conducted to infer the relationship between Tm groups. They were Tm < 55o C vs 55-65o C, < 55o C vs > 65o C, and 55-65o C vs > 65o C (Figure 5). The correlation coefficient (*r*) for the group < 55°C vs 55-65°C, < 55°C vs > 65°C, and 55-65°C vs > 65°C was -0.006, 0.011, and -0.003, respectively (Figure 5). The frequency distribution of *A. thaliana* proteomes with different Tm was depicted in Figure 4. When a correlation plot was drawn to understand the role of Tm, *pI*, and molecular mass (kDa), it was found that molecular mass has a role towards the Tm of *Arabidopsis* proteins with a correlation coefficient *r* = 0.259. However, the *pI* of the *Arabidopsis* proteome is negatively correlated with coefficient *r* = -0.091 (Figure 6).

**Figure 4.**
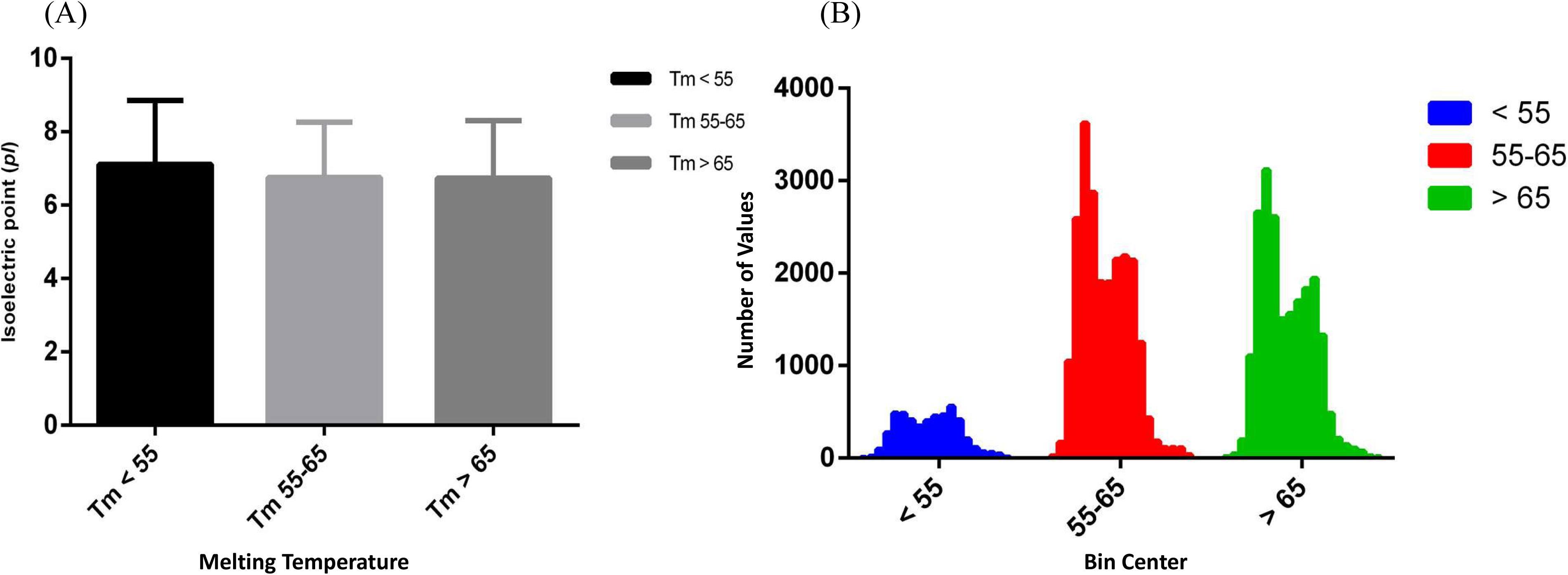
Figure depicting the (A) isoelectric point and (B) frequency of occurrence of isoelectric point of *A. thaliana* proteome in three different Tm groups. The isoelectric point of proteins with Tm < 55°C was found to be the highest in *A. thaliana* proteome.

**Figure 5.**
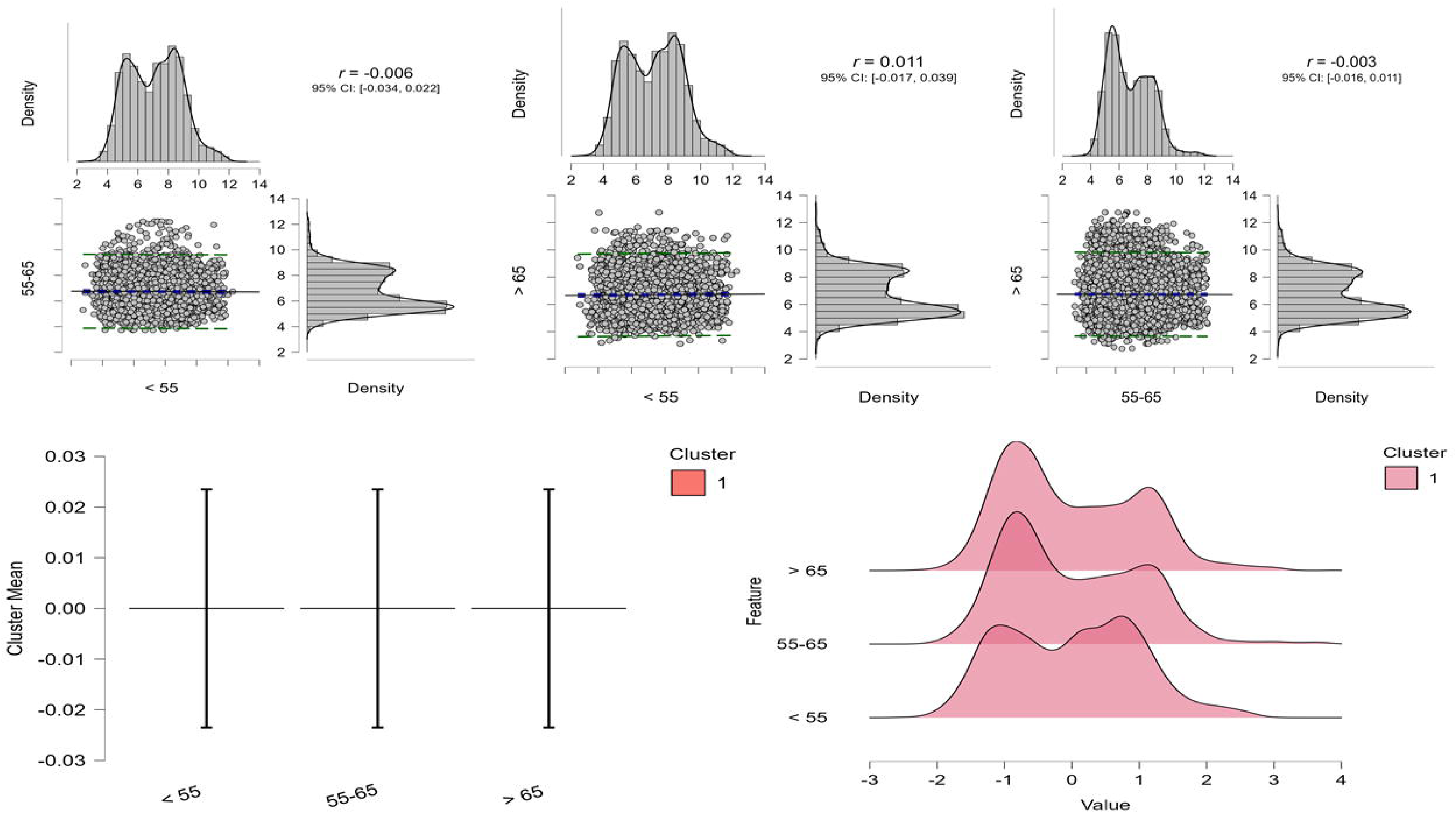
Correlation analysis of isoelectric points of *A. thaliana* proteome within three different Tm groups. The Tm groups were < 55°C, 55-65°C, and > 65°C. The correlation study was carried out with groups < 55°C vs 55-65°C, < 55°C vs > 65°C, and 55-65°C vs > 65°C. Analysis shows, correlation coefficient between the Tm group of < 55° C vs > 65° C was highest (r = 0.011). Pearson’s correlation was used for the correlation study with p < 0.05. The analysis was conducted using JASP software, version 0.19.0.0.

**Figure 6.**
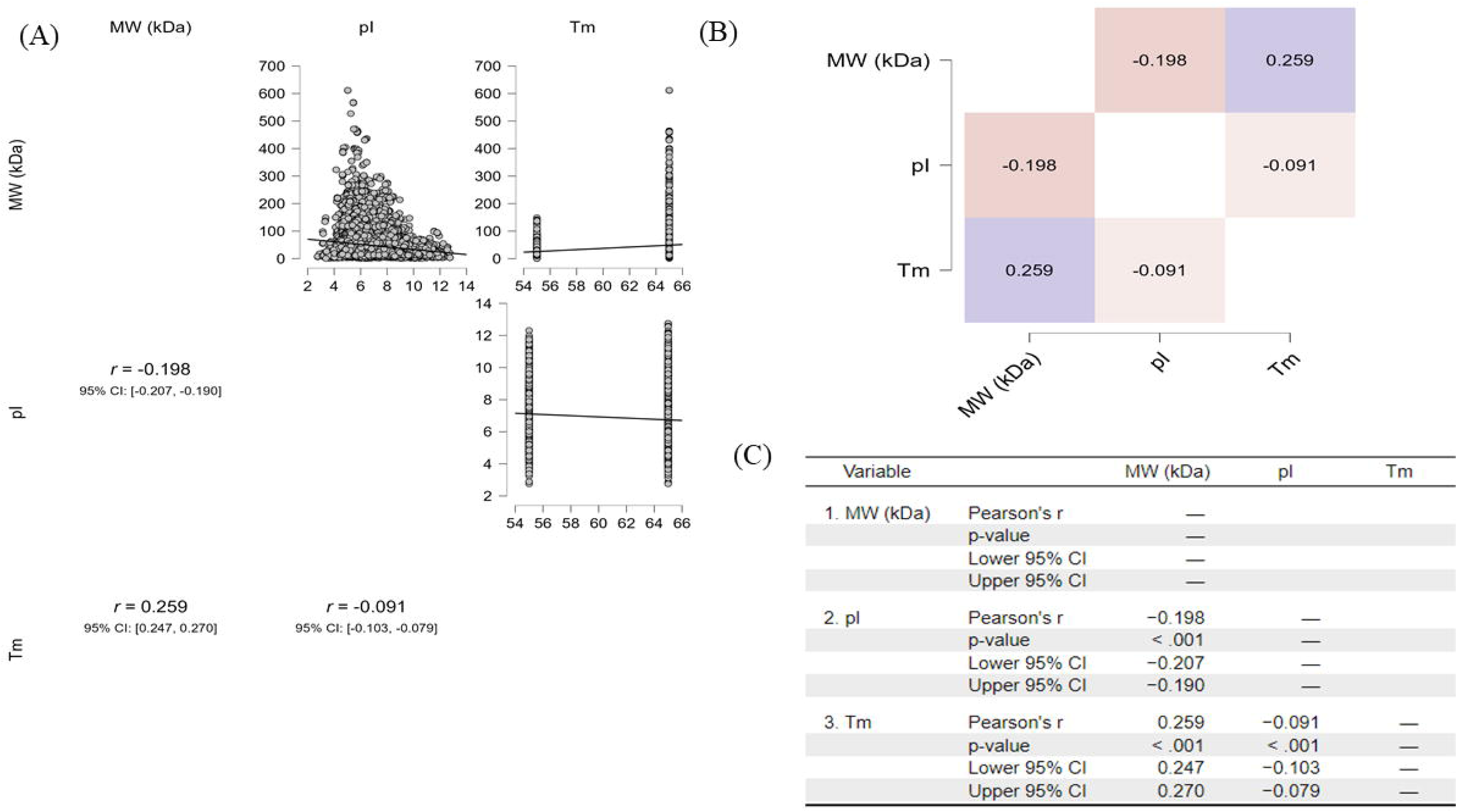
Figure depicting the (A) correlation analysis between the molecular weight, isoelectric point, and Tm of the *A. thaliana* proteome. Figure (B) shows the heat map of the correlation and (C) presents the statistical details of the correlation study. The figure shows positive correlation between Tm and molecular weight (kDa) (*r* = 0.259) while Tm and *pI* (*r* = -0.091) and molecular weight and *pI* (*r* = -0.198) had negative correlation. Pearson’s correlation was used in this analysis with *p* < 0.01. The analysis was conducted using JASP software version 0.19.0.0.

### Machine Learning Approach Showed Molecular Mass has Influence on Protein Tm

To understand the role of molecular mass, isoelectric point, and Tm index in deciding the Tm of *A. thaliana* protein, a machine learning approach was adopted to find their role. We conducted a boosting regression to understand the influence of different variables on Tm. It was found that molecular mass has a relative influence on Tm than the isoelectric point (Figure 7). The study contained 30950 training, 7738 validation, and 9671 test sets. Again, a decision tree regression study was conducted, and it was found that molecular mass (kDa, n = 30950) plays an important role in the Tm of the *Arabidopsis* proteins (Figure 8). Further, a network plot analysis of molecular mass (kDa), Tm, and *pI* was conducted. It was found that kDa and Tm are positively correlated (blue) while kDa and TI are negatively correlated (red) (Figure 8). However, *pI* was only linked to kDa, and it did not show any network with TI and Tm of proteins (Figure 8D).

**Figure 7.**
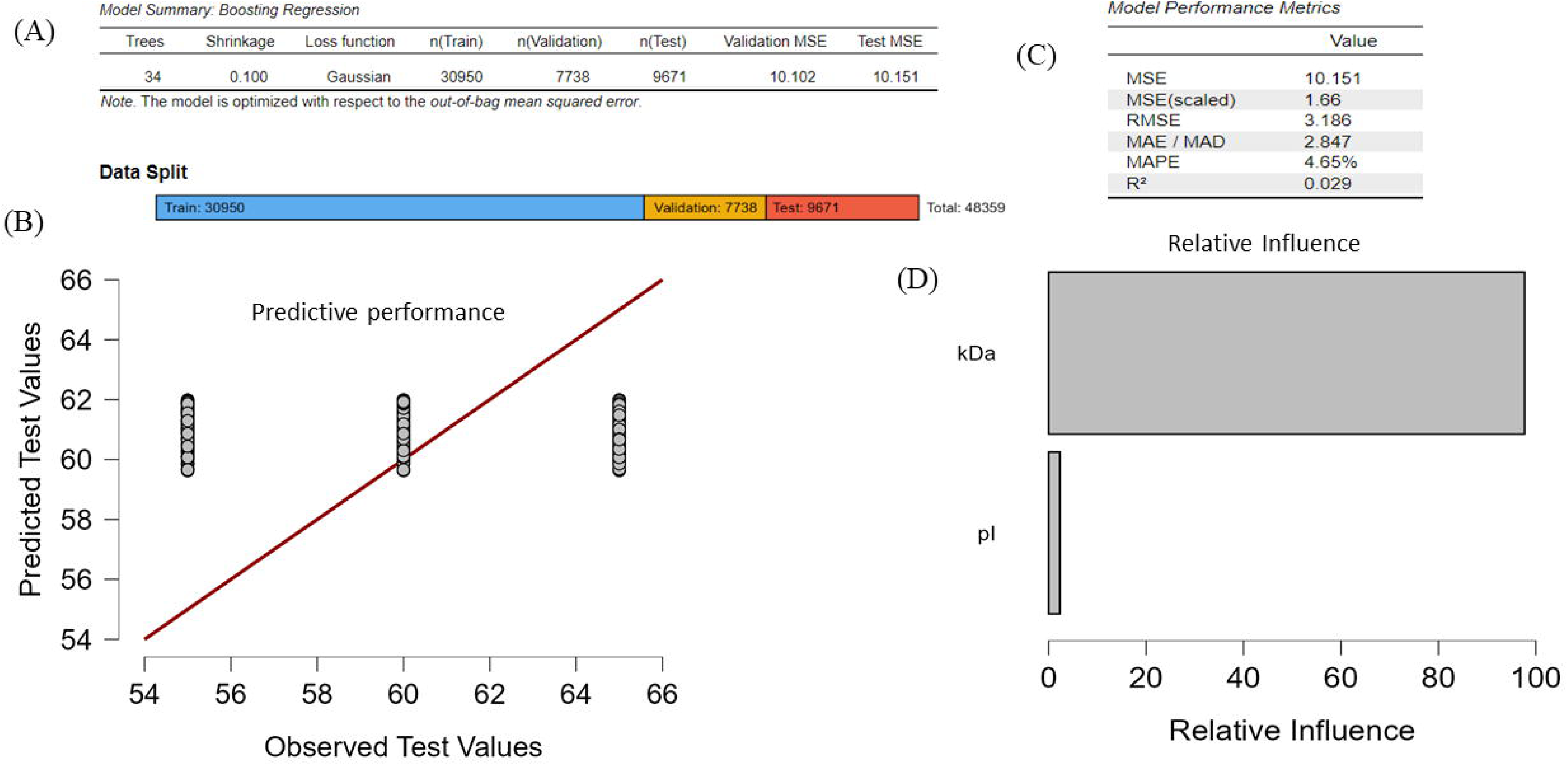
Machine learning analysis of molecular weight (kDa) and isoelectric point to identify their role in deciding the Tm of the *A. thaliana* proteome. Figure depicts (A) boosting regression, (B) data splits, (C) model performance matrics, and (D) relative influence of the data. In the study, there was 30950 training, 7738 validation, and 9671 test set data. The boosting regression model was optimized with respect to out-of-bag mean square error. The analysis revealed the influence of molecular weight (kDa) towards deciding the Tm of *A. thaliana* proteome. The analysis was conducted using JASP software version 0.19.0.0.

**Figure 8.**
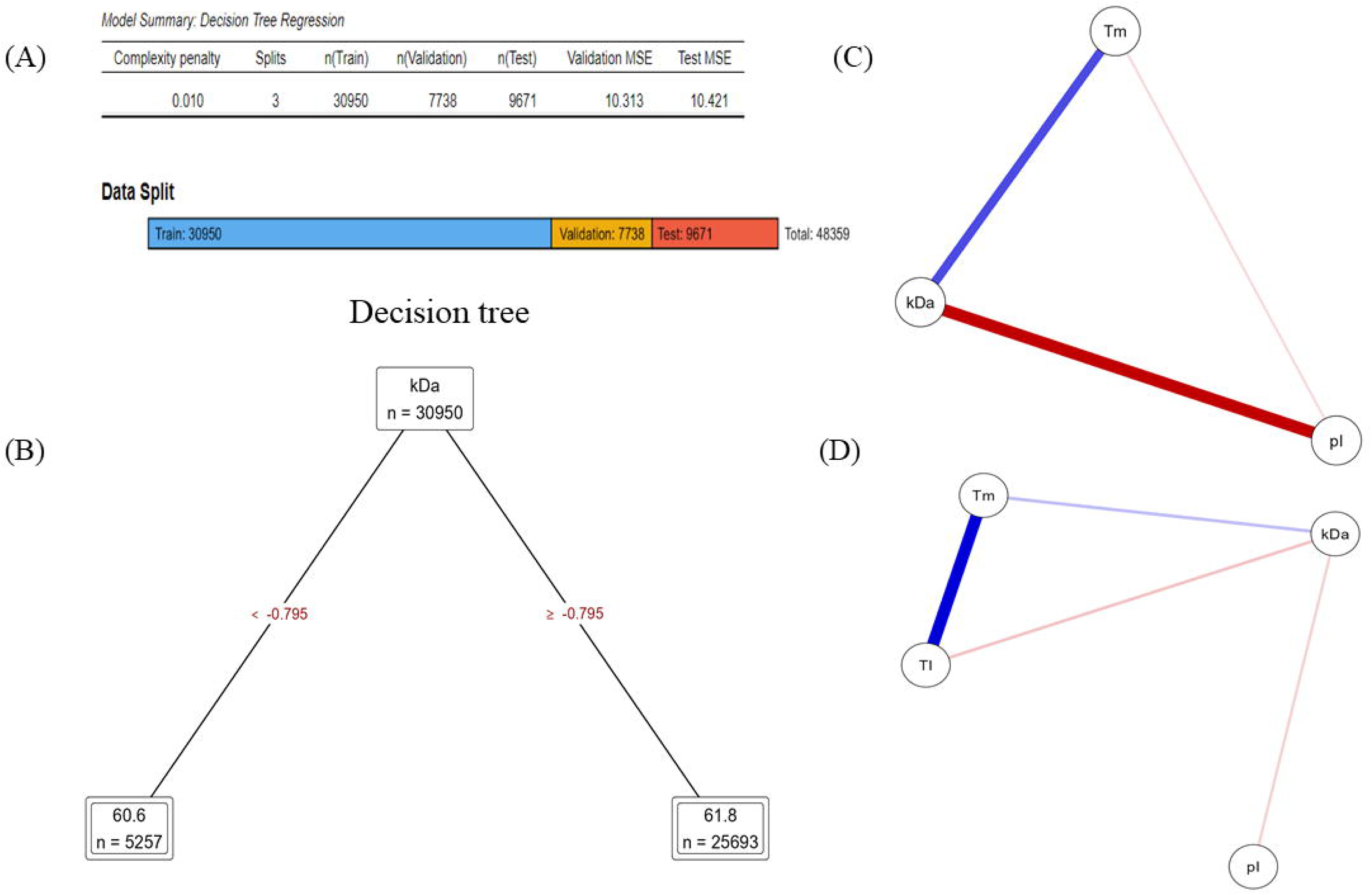
Decision tree regression analysis of molecular weight and isoelectric point of *A. thaliana* proteome. The figure depicts (A) decision tree regression with data split (B) Decision tree (C) network-plot between molecular weight (kDa), Tm, and *pI*, and (D) network-plot between Tm, kDa, *pI*, and TI (Tm index). The analysis contained 30950 training, 7738 validation, and 9671 test set data. The analysis resulted kDa (n = 30950) in the decision tree towards deciding its role in Tm of *A. thaliana* proteome. Network-plot shows, Tm and kDa (Fig. 8C) positively related (blue) while kDa and *pI* negatively related (red). Similarly, there is a positive relationship between Tm and TI and Tm and kDa. While TI and kDa and kDa and *pI* were negatively related.

### High and low TI Encoding Arabidopsis Gene undergone duplication

We picked the top 50 highest TI (Tm > 65°C), middle TI (55-65°C), and lowest TI (< 55°C) encoding CDS sequences of *A. thaliana* proteins and conducted a phylogenetic analysis separately. The phylogenetic tree of *A. thaliana* genes with protein TI < 55°C showed two major clusters (Figure 9A). Although the proteins were from diverse groups, their clustering in the phylogenetic tree was quite smooth. The phylogenetic tree of proteins with Tm < 55°C showed all of the genes were duplicated (Table 3). None of the genes was found to undergo loss, transfer, or codivergence (Figure 9A, Table). The phylogenetic tree of genes with TI group 55-65°C also resulted in two major clusters (Figure 9B), while the phylogenetic tree of genes with TI > 65°C showed three distinct clusters (Figure 9C). However, all the genes of TI < 55°C, 55-65°C, and > 65°C underwent duplication, and none of them were found to undergo loss, transfer, or codivergence (Supplementary Figure 1, Supplementary Figure 2, Supplementary Figure 3, Table 3) during evolution.

**Figure 9.**
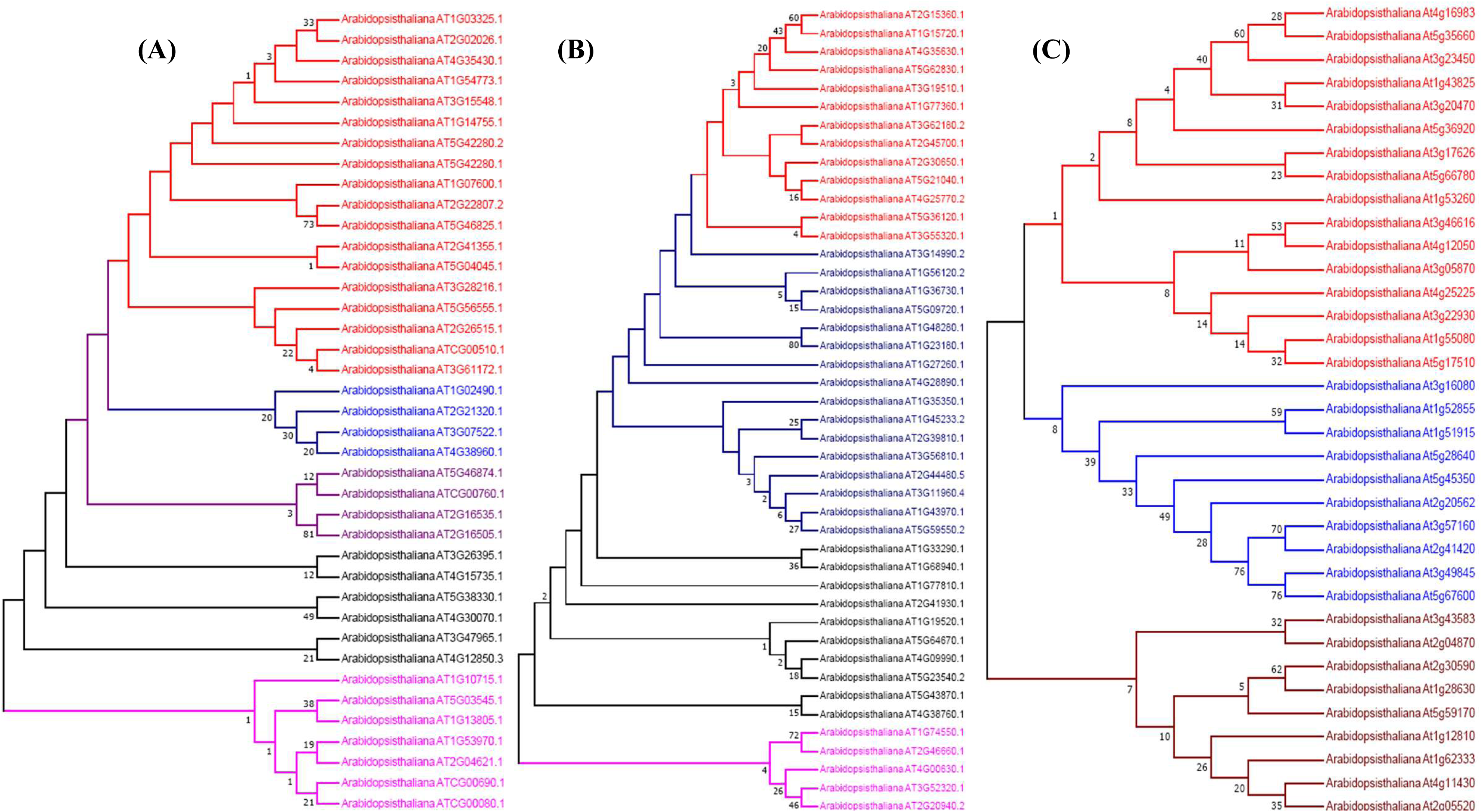
Phylogenetic trees of A. thaliana CDS sequences with different Tm groups. The phylogenetic tree (A) belongs to the CDS of proteins with the Tm < 55°C (B) belongs to the CDS of proteins with the Tm group 55-65°C, and (C) belongs to the CDS of proteins with the Tm > 65°C. Top 50 highest TI containing protein sequences from each Tm group was considered to construct the phylogenetic tree. The CDS sequences of the protein Tm group > 65°C resulted only three major clusters while the CDS sequences of other Tm group resulted multiple clusters. This reflects, the protein sequences of the Tm group with > 65°C were evolutionarily more closer with each other than the other Tm groups. The phylogenetic tree was constructed using MEGA software version 7.

Further, we tried to understand the relative synonymous codon usage (RSCU) of the genes those used for the phylogenetic analysis. We found that the AGA codon coding for the Arg amino acid has the highest RSCU for genes for the Tm group < 55°C (2.33) and 55-65°C (2.26) (Supplementary File 1). However, the highest RSCU for genes for the Tm group > 65°C was found in codon CCA (1.8), which codes for the Pro amino acid. Similarly, the lowest RSCU for the Tm group < 55°C (0.38) and 55-65°C (0.44) group was found in the codon GCG that encodes for the Arg amino acid (Supplementary File 1). However, for Tm group > 65°C, the lowest RSCU was found in codon CCC (0.36), which encodes for the Pro amino acid (Supplementary File 1). In the compositional analysis of nucleotides, the nucleotide composition for the Tm group < 55°C was T = 30.2%, C = 19.5%, A = 28.3%, and G = 21.9%; for Tm group 55-65o C, T = 27.6%, C = 19.8%, A = 28.5% and G = 24.1%; and for Tm group > 65o C, T = 22.3%, C = 23.3%, A = 27.9%, and G = 26.4%.

### Nucleotide Position in Codon Explain genetic variation and adaptive role for stress tolerance

We analyzed the nucleotide position in the codon of the studied genes to understand and evaluate their roles towards shaping the protein structure, function, and evolution. We analyzed the nucleotide position of the codons of different Tm groups. In the Tm group < 55°C, at the first position, the abundance of T (31) was followed by A (29.5), G (22), and C (18.2) (Figure 10); at the 2nd position the abundance of T (31) was followed by A (27.2), G (24.6), and C (17.7) (Figure 10); at the 3rd position the abundance of T (32.04) was followed by A (27.6), C (20.4), and G (19.9) (Figure 10). In the Tm group 55-65°C, at the 1st position, the abundance of nucleotide A (29.2) was followed by T (28), G (23.3), and C (19.8) (Figure 10); at the 2nd position, the abundance of nucleotide A (28.1) was followed by T (27), G (24.7), and C (20.4) (Figure 10); at the 3rd position, the abundance of nucleotide A (28.3) was followed by T (27), G (24.5), and C (20.2) (Figure 10). In the Tm group > 65°C, at the 1st position, the abundance of nucleotide A (26.7) followed by C (25.1), G (24.1), and T (23) (Figure 10); at the 2nd position, the abundance of nucleotide G (28.4) followed by A (26.9), C (22.5), and T (22) (Figure 10); at the 3rd position, the abundance of nucleotide A (26.9) followed by T (25), C (24.3), and G (24.2) (Figure 10).

**Figure 10.**
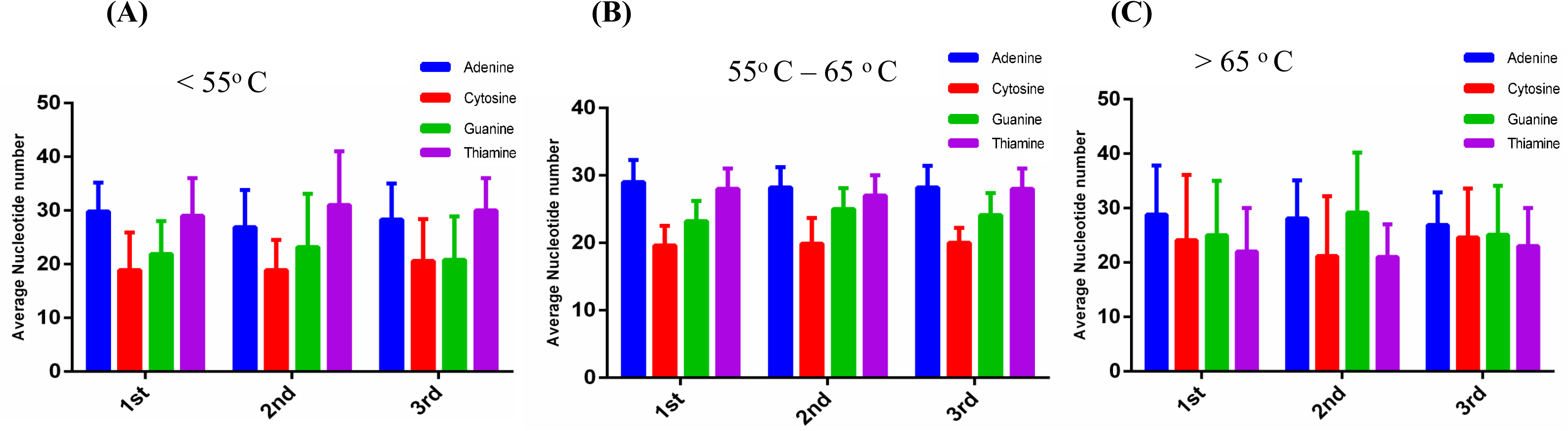
The figure illustrates the nucleotides position and variation in the codons in different position. The positions of the nucleotides adenine (A), cytosine (C), guanine (G), and Thiamine (T) at 1^st^, 2^nd^, and 3^rd^ was studied to understand their preferences in the codon. The nucleotide positions in the codon was studied with three different melting temperature groups (A) < 55°C, (B) 55-65°C, and (C) > 65°C.

When we analyzed the nucleotide position by grouping them with the Tm group (e.g., A nucleotide at the 1st position for the Tm group < 55°C, 55-65°C, and > 65°C). We found the frequency of A nucleotide (28.46) at the 1st position was highest for all three Tm groups, followed by T at the 3rd position (27.87), A at the 3rd position (27.62), and A at the 2nd position (27.38) (Figure 11). The frequency of C at the 2nd position was the lowest (20.18), followed by C at the 1st (21.03) and 3rd positions (21.63) (Figure 11). When we analyzed the frequency of nucleotide variations for all three Tm groups at the 1st, 2nd, and 3rd positions, we found the frequency of nucleotide variance was highest in T at the 2nd position (17.34), followed by T at the 3rd (14.59), T at the 1st (13.83), and C at the 1st (13.06) positions (Figure 11). The lowest nucleotide variance was recorded for A at the 2nd position (0.37), followed by A at the 3rd position (0.49) and G at the 1st position (1.81). Further, a cluster analysis revealed the frequency of nucleotide positions for the Tm group < 55°C and the 55-65°C group fall close to each other, while the nucleotide position of the Tm group > 65°C grouped separately.

**Figure 11.**
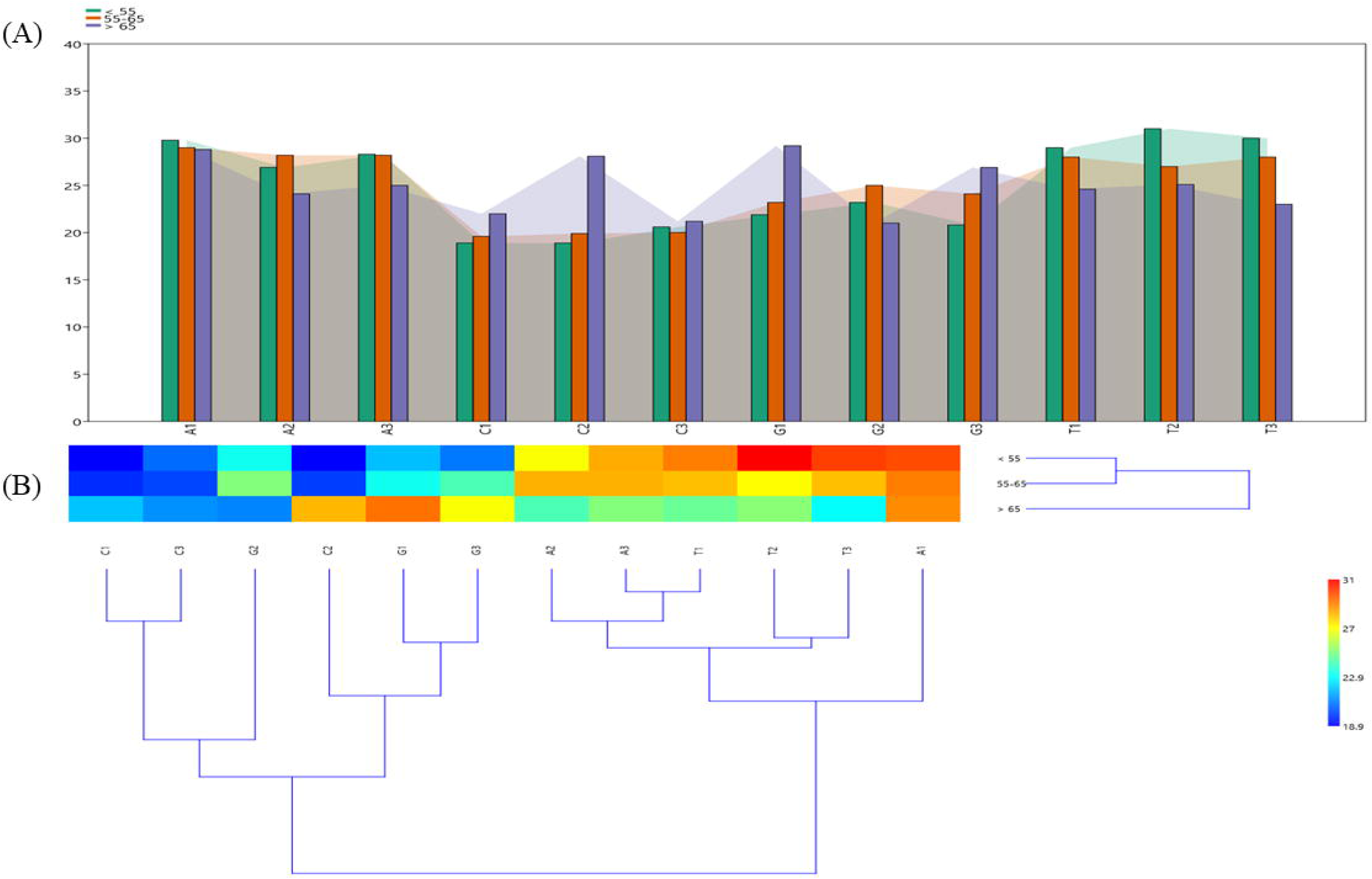
Melting temperature specific nucleotide position of codons in A. thaliana CDS sequences. Nucleotide position of three different Tm groups were clubbed together to understand their frequency. There is a significant variation of Cytosine nucleotide at 2^nd^ position in the Tm > 65°C. Similarly, the frequency of Guanine nucleotide was quite higher in Tm > 65°C compared to other Tm groups. Cluster analysis revealed, the nucleotide positions with Tm groups < 55°C, & 55-65°C grouped together while nucleotides position of Tm group > 65°C fall separately. This signifies there is a significant variation in nucleotide position in the codon of the protein having Tm > 65°C.

### Relative Synonymous Codon Usage (RSCU) of A. thaliana Genes

We conducted a study to understand the relative synonymous codon usage of genes of the top 50 *A. thaliana* proteins from three Tm groups. We found the RSCU of codon AGA (R) was highest (2.33) in the Tm group < 55°C and 55-65°C (2.26). For the Tm group > 65°C, the RSCU of CCA (P) was highest (1.8) (Supplementary file). The lowest RSCU was found in CCC (P) (0.36) in the Tm group > 65°C, followed by CGC (R) (0.38) in the Tm group < 55°C and 0.44 in the Tm group 55-65°C (Supplementary File). The second highest RSCU was found in the stop codon UGA in the Tm group < 55°C and 55-65°C. From the 64 codons, at least 27 codons contained RSCU of > 1 in Tm group < 55 C while 34 codons contained RSCU of < 1. Only three codons (AUG (M), CCU (P), and UGG (W)) of the Tm group < 55°C contained an RSCU value of 1. For the Tm group 55-65°C, 28 codons contained RSCU of > 1, while 32 codons contained RSCU < 1. Four codons (CCU (P), UGG (W), CGR (R), and ACU (T)) in the Tm group 55-65°C were found to contain an RSCU value of 1. For the Tm group > 65°C, 31 codons contained RSCU > 1, and a similar number of codons were found to contain RSCU < 1. Only two codons, AUG (M) and UGG (W), contained an RSCU value of 1 in the Tm group > 65°C.

### Tm Based Codon Context of Arabidopsis CDS

Codon usage preferred the use of codons; the “codon context” explains the sequential presence of codon pairs in a gene. To understand the codon pairs in *Arabidopsis* CDS with protein Tm groups of < 55°C, 55-65°C, and > 65°C, we conducted codon pair analysis. For CDS of protein Tm group < 55°C, ATG ATG was the preferred codon pair and found in the highest number (37), followed by TGT GAT (13), ATG GAA (10), CTT TGC (9), and AAA TGT (8). For the Tm group 55-65°C, the GAA GAT and GAG AAG (30) codon pairs were found in the highest number, followed by GAT GAA (25), GAT GAT (25), AAA GAG (24), AAA GAT (23), GAA GAA (23), AAA GAA (21), and others. For the Tm group > 65°C, the CAA CAA (58) codon pair was found in the highest number, followed by GGA GGA (49), CCA CCA (44), GGT GGA (38), GGT GGT (32), and TAT CCT (31).

## Discussion

The proteins are developed to perform their function within a specific temperature range. However, the stability of a plant proteome is directly proportional to the survival rate of the plants under the heat stress. Heat stress can disrupt the cellular and extracellular proteins, leading to the disruption in the cellular homeostasis, physiological process, and metabolism. The inactivation of important and functional proteins can lead to compromised enzyme activities that will impact the enzymatic pathways in the plant. Therefore, understanding the protein stability for temperature resistance is an important parameter to study the heat stress in plants. Thermal stability of protein, referred to as melting temperature (Tm), provides the strength to the protein to cope with extreme temperature stress. The melting temperature of the protein is defined as the temperature at which the protein undergoes a transition from native, folded form to unfolded form under equilibrium conditions. The proteins having high Tm will be less prone to denaturation and structural instability, leading to high cellular integrity and function under adverse heat conditions. It can be well speculated that proteins with high Tm can better adapt to perform their function under high-temperature conditions, thus making Tm a critical thermodynamic parameter for understanding heat resistance in plants. Due to the denaturation by high temperature, protein will lose its functional conformation, leading to its inactivation. Therefore, it is important to understand the thermodynamic properties of proteins, and hence we have conducted a study to deduce the melting temperature (Tm) of the entire *Arabidopsis thaliana* proteome. For a better understanding, protein Tm was grouped into three groups with temperature ranges < 55°C, 55-65°C, and > 65°C. However, the Tm of the protein relies on several factors, including amino acid composition, structure, and environmental factors, including pH, ionic strength, and the presence of stabilising and destabilizing factors. From these factors, amino acid composition is one of the major parameters that might be controlling the Tm and stability of the protein. Therefore, we conducted an in-depth analysis of the amino acid composition of the *Arabidopsis thaliana* proteome. We found 20640 proteins (including splice variants) of *A. thaliana* fall in Tm range > 65°C, 22826 proteins fall in the Tm range 55°C– 65°C, and 4893 proteins fall in the Tm range < 55°C. From this study, it is evident that *A. thaliana* has developed more proteins with Tm > 65°C. This shows the plant has developed proteins to withstand high temperatures. Considering the evolutionary consequences, the earth has passed two ice ages and subsequently gained temperature, and now we are facing global warming. During this evolutionary process, plants have developed their protein machinery systems and encoded more proteins that can withstand high-temperature stress. This evolving process of rising temperatures might continue in the future, and hence we need to find ways to identify important heat stress-resistant proteins to tackle the global warming problem. Therefore, finding the heat-resistant proteins in plants is of paramount interest to understand their heat-resistant mechanism.

To have a better understanding of high and low Tm proteins, we studied the amino acid composition of the *A. thaliana* proteome. It was found that the amino acid composition of Ala, Asp, Glu, Gly, Lys, Leu, Gln, and Val increased with an increase in the Tm of the proteins, while the composition of Cys, Phe, His, Ile, Asn, Pro, Arg, Ser, Thr, Trp, and Tyr decreased with an increase in the Tm of the protein (Figure 1, Table 2). This linearity of increase and decrease in amino acid composition reflects their role in determining the Tm of the proteins. Further, we calculated the molecular mass and isoelectric point of individual proteins of *A. thaliana* and studied their role towards the Tm. The average molecular mass of *A. thaliana* for protein Tm group < 55°C was 25.61 kDa, for Tm group 55-65°C it was 52.48 kDa, and for Tm group > 65°C it was 48.92 kDa (Figure 2). The molecular mass of proteins with Tm < 55°C was quite low compared to the Tm group 55-65°C and > 65°C (Figure 2). This gives a hint that proteins with low molecular mass possess low Tm. A correlation analysis within the molecular mass group showed proteins with Tm < 55°C marginally correlate with the proteins with Tm 55-65°C (Figure 3). Analysis of the isoelectric point of *A. thaliana* protein revealed that proteins with Tm < 55°C possess high *pI* (7.22), while the *pI* of proteins of the Tm group 55-65°C was 6.75 and the *pI* of the Tm group > 65°C was 6.74 (Figure 5). This shows that high *pI* protein tends towards low Tm in *A. thaliana*. The trend of *pI* of the proteins for three different groups was in the decreasing order from Tm < 55°C to > 65°C. However, the *pI* difference was not significant enough. Similarly, there were no such detectable correlations in the isoelectric point of proteins found in three different Tm groups (Figure 6). Further, a correlation study was conducted using molecular weight, isoelectric point, and Tm as input parameters, and the result showed a positive correlation between molecular weight and Tm (Figure 8). The relative influence of molecular weight (kDa) on the Tm of *A. thaliana* protein was further confirmed using a machine learning approach (Figure 9). Boosting regression analysis revealed the relative influence of molecular weight (kDa) on the melting temperature of the proteins. Further, the validation role of molecular weight (kDa) towards the melting temperature of protein was observed in the decision tree regression analysis and network plot study (Figure 10).

A study regarding the melting point analysis was reported by Jarzab et al. (2020), where they studied 48000 protein sequences from humans and archaea and covered 13 species [82]. They reported that protein sequence, composition, and size affect the thermal stability of proteins in prokaryotes and eukaryotes [82]. They used the species that lived in the 40°C to 70°C temperature range. It is well known that the enzyme DNA polymerase from the bacteria *Thermus aquaticus* is highly resistant to heat, having a half-life period of 40 minutes at 95°C. The study reported that RNA polymerase of *Thermus aquaticus* remains unaltered at 80°C and helps in synthesising tRNA transcripts [83]. In our study, we found the mediator protein of the RNA polymerase II transcription subunit-like protein that falls in the Tm > 65°C temperature range has the highest Tm index (TI = 9.60). This shows the mediator protein of RNA polymerase II might be contributing towards the thermal stability of the RNA polymerase. Further, proline-rich, cysteine-rich, calmodulin-like proteins, and transmembrane proteins were found to contain higher Tm index (Table 1). This shows that proteins involved in transmembrane activity, calcium signaling, and proline metabolism are associated with heat resistance in *A. thaliana*. Tang et al. reported the melting temperature of 95°C in legume protein edestin and found that the melting temperature was unaffected in the presence of 20-40 µM sodium dodecyl sulphate [84]. Gliadin protein functions in a temperature range of 70-115°C, and as the temperature rises to 135°C, gliadin protein softens much better [85]. The thermal stability of plant H2-ferritin protein (soybean) was reported to be quite high. When H2-ferritin protein is caged with extra peptides, it denatures at 106o C [86]. Miller et al. (2013) reported the thermal stability of ribulose-1,5-bisphosphate (Rubisco) and measured the Tm of native, ancestral, and variant proteins from *Synechococcus*. They found that OH28 purified Rubisco exhibited greater stability at 79.5°C than the less thermotolerant strain (72.3-73.6°C) [87]. This led them to conclude that thermostable Rubisco enzyme diverged from the ancestor of *Synechococcus* [87]. Further, codon evolution revealed the substitution of the Ala amino acid by isoleucine in the OH28 strain, suggesting a positive selection in evolution [87]. To understand the evolutionary aspects of the high Tm, low, and medium Tm proteins, we conducted gene duplication and loss analysis. We found that all the studied genes of three Tm groups had undergone duplication, and none of them were found to undergo transfer, codivergence, or losses (Table 3). Gene duplication is an important event as it accumulates mutations over time without affecting the function of the original gene, thus bringing the new function and characteristics. The duplication of the genes with high, low, and medium Tm genes reflects their tendency towards the innovation of new functions. In gene duplication, one gene can continue to exhibit its original function while allowing others to evolve new functions, leading to the adaptability of the species to adverse temperature conditions.

The RSCU value of 1 indicates a codon is used in expected frequency, and < 1 indicates the codons are used quite less frequently. However, RSCU of > 1 shows, codons are used more frequently than expected. For the Tm group < 55°C, 27 codons contained RSCU > 1, and 32 codons contained RSCU < 1; for the Tm group 55-65°C, 28 codons contained RSCU > 1, and 32 codons contained RSCU < 1; for the Tm group > 65°C, 31 codons contained RSCU > 1, and 31 codons contained RSCU < 1. From them, the Tm group < 55°C and 55-65°C, 27 and 28 codons, respectively, contained codons with an RSCU of > 1, while 32 codons contained an RSCU of < 1. The result depicts RSCU of value > 1 increased with the increase in the Tm of the proteins, while the RSCU of value < 1 decreased with the increase in the Tm of the proteins. In the Tm group at < 55°C, only three codons had an RSCU of 1; in the Tm group at 55-65°C, only four codons contained an RSCU of 1, and in the Tm group at > 65°C, two codons were found to contain an RSCU of 1. The RSCU of codons having value 1 for all the Tm groups were AUG (M) and UGG (W) (Supplementary File). For the Tm group < 55°C, unique codon with RSCU value 1 was CCU (P) while for Tm group 55-65°C unique codons with RSCU 1 were CUC (L) and AGC (S). The second highest RSCU was found in codon UGA for the Tm group < 55°C and 55-65°C. The UGA codon also plays an important role in encoding selenocysteine amino acid in the protein. This analysis revealed that *A. thaliana* encodes differential RSCU according to the Tm groups, reflecting the role of codons in encoding proteins with different melting temperatures. The codon usage bias in the different Tm groups was quite evident. The RSCU keeps the protein sequence intact while altering their influences on gene expression, protein translation, and stress responses. The RSCU bias is influenced by several factors, including natural selection, mutation pressure, translation efficiency, and gene expression levels. The presence of higher RSCU in codon AGA that encode for Arg amino acid in the Tm group < 55°C and 55-65°C reflects its role in lower to moderate temperature while the higher frequency of codon CCA that encodes for Pro amino acid in the Tm group > 65°C shows CCA codon is optimized for translational accuracy under stress condition. The RSCU study is a vital tool that helps to understand plant adaptation to environmental stress conditions by influencing protein translation efficiency and protein folding. However, the codon usage bias may vary in tissue-specific expression and developmental stages of plants.

The position of nucleotide at the 1st place for all the Tm groups showed the frequency of nucleotide A was the highest. The highest frequency of nucleotide A across the category suggests its vital roles in codons for encoding amino acids critical for protein function. Its higher frequency at Tm > 65°C shows its importance in protein function towards high-stress adaptations. The G nucleotide position has quite high variability, suggesting its role in genetic diversity and potential for adaptive evolution. Nucleotide C is more prone to mutation without severely impacting the protein function, thus providing a buffer for evolutionary changes. Similarly, nucleotide T showed higher consistency, indicating its role in encoding essential amino acids. Overall, it suggests that nucleotides for Tm < 55°C were less variable, for Tm 55-65°C, intermediate variable, and the Tm > 65°C was highly variable. The high variability of nucleotides indicates genomic regions with higher potential for evolutionary innovation, contributing to the genomic diversity and adaptation. The presence of higher nucleotide variability in the Tm group > 65°C indicates *A. thaliana* plants are inclining more towards enhancing the Tm of their protein for stress adaptation while keeping the conserved genomic architecture in the Tm group < 55°C. A comparative less variable nucleotides in different position of codons reflect their conserved structure and their association with protein of the Tm group < 55o C reflects the plants have evolved from cold climatic conditions and now diverged its genomic pool towards stress adaptation. The nucleotide positions in the groups A1, A2, A3, C1, C2, C3, G1, G2, G3, T1, T2, and T3 with their variation reflect a delicate balance between the conservation and stress adaptation in the evolution of genes.

## Declaration

### Ethics approval and consent to participate

Not applicable

### Consent for publication

Not applicable

### Availability of data and materials

All the data associated with the study is presented in the article.

### Author contributions

KMM: identified the melting temperature of the proteins, revised the manuscript; TKM: conceived the idea, analyzed the data, drafted and revised the manuscript

## Acknowledgement

The authors would like to express their sincere thanks to Indian School Nizwa, Oman, and Medi-Caps International School, Indore, Madhya Pradesh, India for their support and encouragement to conduct this research. The authors would also like to extend their sincere thanks to Mrs. Asma Khan, Indian School Nizwa, Oman for her extensive encouragement to author Karan Martens Mohanta to conduct the research.

## Supplementary Materials

**Supplementary Figure 1.** Phylogenetic tree of A. thaliana CDS of < 55° C Tm group. The evolutionary history was inferred by using the Maximum Likelihood method based on the Tamura-Nei model. The tree with the highest log likelihood (-12492.72) is shown. The percentage of trees in which the associated taxa clustered together is shown next to the branches. Initial tree(s) for the heuristic search were obtained automatically by applying Neighbor-Join and BioNJ algorithms to a matrix of pairwise distances estimated using the Maximum Composite Likelihood (MCL) approach, and then selecting the topology with superior log likelihood value. A discrete Gamma distribution was used to model evolutionary rate differences among sites (5 categories (+G, parameter = 7.2022)). The analysis involved 39 nucleotide sequences. Codon positions included were 1st+2nd+3rd+Noncoding. There were a total of 2093 positions in the final dataset.

**Supplementary Figure 2.** The evolutionary history was inferred by using the Maximum Likelihood method based on the Tamura-Nei model. The tree with the highest log likelihood (-95417.24) is shown. The percentage of trees in which the associated taxa clustered together is shown next to the branches. Initial tree(s) for the heuristic search were obtained automatically by applying Neighbor-Join and BioNJ algorithms to a matrix of pairwise distances estimated using the Maximum Composite Likelihood (MCL) approach, and then selecting the topology with superior log likelihood value. A discrete Gamma distribution was used to model evolutionary rate differences among sites (5 categories (+G, parameter = 6.8373)). The tree is drawn to scale, with branch lengths measured in the number of substitutions per site. The analysis involved 44 nucleotide sequences. Codon positions included were 1st+2nd+3rd+Noncoding. There was a total of 6825 positions in the final dataset.

**Supplementary Figure 3.** The evolutionary history was inferred by using the Maximum Likelihood method based on the Tamura-Nei model. The tree with the highest log likelihood (-23319.50) is shown. The percentage of trees in which the associated taxa clustered together is shown next to the branches. Initial tree(s) for the heuristic search were obtained automatically by applying Neighbor-Join and BioNJ algorithms to a matrix of pairwise distances estimated using the Maximum Composite Likelihood (MCL) approach, and then selecting the topology with superior log likelihood value. A discrete Gamma distribution was used to model evolutionary rate differences among sites (5 categories (+G, parameter = 10.5572)). The analysis involved 35 nucleotide sequences. Codon positions included were 1st+2nd+3rd+Noncoding. There were a total of 1757 positions in the final dataset.

**Supplementary File 1.** Relative synonymous codon usage of Arabidopsis thaliana genes with Tm group < 55°C, 55-65°C and < 55°C, & > 65°C.

