## Supplementary Figure 3 for "Meltome Atlas of *Arabidopsis thaliana* Proteome: A Melting Temperature-Based Identification of Heat & Cold Resistant Proteins"

**Supplementary Figure 3**. The evolutionary history was inferred by using the Maximum Likelihood method based on the Tamura-Nei model. The tree with the highest log likelihood (-23319.50) is shown. The percentage of trees in which the associated taxa clustered together is shown next to the branches. Initial tree(s) for the heuristic search were obtained automatically by applying Neighbor-Join and BioNJ algorithms to a matrix of pairwise distances estimated using the Maximum Composite Likelihood (MCL) approach, and then selecting the topology with superior log likelihood value. A discrete Gamma distribution was used to model evolutionary rate differences among sites (5 categories (+G, parameter = 10.5572)). The analysis involved 35 nucleotide sequences. Codon positions included were 1st+2nd+3rd+Noncoding. There were a total of 1757 positions in the final dataset.

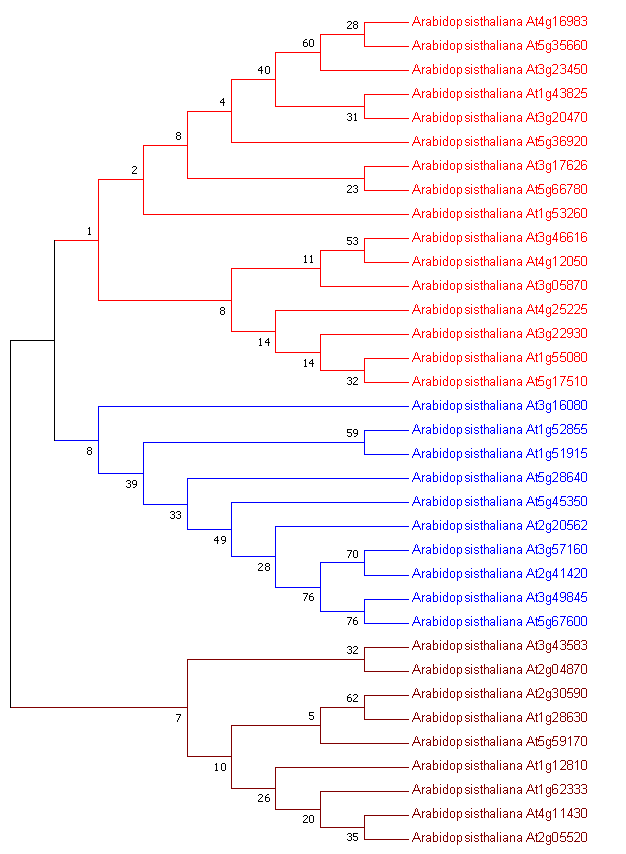
